# Vibrational Spectroscopy Identifies Myocardial Chemical Modifications in Heart Failure with Preserved Ejection Fraction

**DOI:** 10.1101/2023.03.20.533577

**Authors:** Leonardo Pioppi, Reza Parvan, Alan Samrend, Gustavo Justo da Silva, Marco Paolantoni, Paola Sassi, Alessandro Cataliotti

## Abstract

**Background:** Vibrational spectroscopy can be a valuable tool to monitor the markers of cardiovascular diseases. In the present work, we explored the vibrational spectroscopy characteristics of the cardiac tissue in an experimental model of heart failure with preserved ejection fraction (HFpEF).

**Methods:** We used the Fourier-transform infrared (FTIR) and Raman micro-spectroscopic techniques to provide complementary and objective tools for the histological assessment of heart tissues from an animal model of HFpEF. A new sampling technique was adopted (tissue print on a CaF2 disk) to characterize the extracellular matrix.

**Results:** Several spectroscopic markers (lipids, carbohydrates, and glutamate bands) were recognized in the cardiac ventricles as due to the comorbidities associated with the pathology, such as obesity and diabetes. Besides, abnormal collagen cross-linking and a decrease in tryptophan content were observed and related to the stiffening of ventricles and to the inflammatory state which is a favorable condition for HFpEF.

**Conclusions:** By the analyses of tissues and tissue prints, FTIR and Raman techniques were shown to be highly sensitive and selective in detecting changes in the chemistry of the heart in experimental HFpEF and its related comorbidities.

## 1. Introduction

It is widely accepted the idea that physiological and pathological conditions can be detected by probing the optical properties of cells and tissues to assess changes in morphology, mechanics, and biochemistry^1^. Although histopathology and immunohistochemistry are currently the gold standards for determining the pathophysiological status of biological samples, the need for new diagnostic tools arises to overcome their limitations^2^. Histological methods rely exclusively on ex-vivo biopsies, which require time-consuming manipulation steps thus causing delayed results^2^. It is necessary to use many chemicals for fixation, inclusion, and staining procedures; these can induce physicochemical changes that significantly alter the structure and composition of the biological specimen of interest, increasing the risk of contamination^3^. Additionally, staining a tissue section can only be used to monitor a few components at once; a complete evaluation requires serial staining and sectioning steps, resulting in bigger sample sizes and longer diagnostic times.

Optical techniques, and specifically vibrational spectroscopies based on infrared (IR) absorption or Raman scattering, offer an alternative to histopathology with some advantages. Besides being non-invasive, non-destructive, and label-free, the Fourier Transformed InfraRed (FTIR) and Raman spectra rely on the total biochemical composition of a sample, revealing the fingerprints of multiple substances in a single measurement^4^. Vibrational spectra can provide useful information not only about the chemical composition of a biological sample but also about its structural properties. This high sensitivity to monitor biochemical and structural changes likely allows to reveal the onset of pathologies even before the morphological and systemic manifestation of the disease^4,5^.

The application of FTIR and Raman spectroscopy to the diagnosis of cardiovascular diseases has been poorly investigated^6–9^. Recent research from Tombolesi et al.^4^ has shown the advantages of the combined use of these techniques in the characterization of cardiac tissue and early diagnosis of heart failure. Heart failure (HF) is a complex clinical syndrome resulting from structural and functional impairment of ventricular blood filling or ejection. There are currently about 64 million people suffering from HF worldwide, and the number continues to rise^10^. HF with preserved ejection fraction (HFpEF) is a medical condition characterized by cardiac remodeling that, at advanced stage, evolves towards concentric hypertrophy, induces left ventricular (LV) wall thickening and changes in the volume and mass of the LV. Studies show that more than half of all patients with HF are affected by HFpEF^11,12^, which is insidious to recognize due to the plethora of risk factors and comorbidities associated with the pathology. The comorbidities accompanying HFpEF are essentially non-cardiac conditions, such as obesity, diabetes mellitus, hypertension, anemia, chronic obstructive pulmonary diseases, and chronic kidney diseases. It is a complex scenario, in which it is important to recognize the different chemical alterations, secondary to comorbidities, that lead to structural and functional abnormalities of the heart.

In this work, we used both FTIR and Raman spectroscopic methods to investigate heart tissues from lean and obese ZSF1 rats which are an established model of HFpEF with combined comorbidities. Changes in the average biochemical composition of the tissue from obese compared to lean rats were related to the advanced stage of HFpEF and cardiac remodeling observed by echocardiography. We monitored the effect of obesity and diabetes as comorbidities, in this respect we compared spectroscopic results with biochemical and histochemical data. To assess the properties of the extracellular matrix (ECM), we analyzed the tissue print on a substrate. This procedure was previously used for the identification of ECM in kidney cancer^13^ and was never applied to the heart. We employed this approach to verify whether the analysis of ECM, other than the whole tissue, provides spectroscopic markers of HFpEF. Once recognized the spectral markers of the pathology, IR and Raman techniques were used to monitor the early stage of the pathology, thus overcoming the limitations of conventional diagnostic tools.

## 2. Materials and Methods

### 2.1 Study protocol

The study conformed to the regulations of our laboratory animal facility (Comparative Medicine, UIO, Norway). The protocol was approved by the Norwegian Food Safety Authority committee (Mattilsynet) for animal research (FOTS protocol number 15886). Four-week-old male ZSF1 Ob) and lean (Ln) were purchased from Charles River Laboratories (USA). Rats were housed in a room with a 12/12-hour light cycle, temperature of 21°C, and humidity of 55%. Rats were maintained on regular chow, water and food were provided ad libitum for the entire course of the study. At the age of 25 weeks, when obese rats developed HFpEF, were euthanized according to American Veterinary Medical Association (AVMA) Guidelines for the Euthanasia of Animals (2020) via deep anesthesia (5% isoflurane), exsanguination and organ excision. Due to obvious phenotype (Ob vs Ln) blindness was not possible for *in vivo* assessments. However, blinding was performed for all the *ex vivo* assessments.

### 2.2 Ultrasounds

Cardiac function was assessed the day before sacrifice by transthoracic echocardiography using a VEVO 3100 high-resolution *in vivo* imaging system from VisualSonics (Amsterdam, NL). Briefly, animals were maintained under anesthesia (1.5-2% isoflurane mixed with oxygen) on a pre-warmed ECG transducer pad with body temperature and ECG monitored. Measurements were made with an MS250 transducer, frequency set at 20 MHz. B-mode measurements in the parasternal long axis view were obtained to assess the function and dimension of the left ventricle (LV). LVEF was calculated as 100 * ((LV Vol;d – LV Vol;s) / LV Vol;d). E and A waves in LV filling velocities were assessed via pulsed-wave Doppler in the parasternal long axis view.

### 2.3 Biochemical assessments

Samples were sent in one batch for a one-kit analysis of blood using an enzymatic method on an automatic biochemical analyzer (Hitachi 7080, Tokyo, Japan). Glucose, cholesterol, and triglycerides were measured in serum.

### 2.4 Blood pressure and hemodynamic measurements

Before sacrifice, animals were placed over a heating platform (preheated to 33 to 35°C), intubated and kept under anesthesia with a mix of 2.5% isoflurane and 97.5% oxygen using the VentElite small animal ventilator (Harvard Apparatus). Invasive measurements were done using Powerlab/Millar modules (ADInstruments) and the SPR-838 rat pressure-volume catheter. Via closed chest approach, arterial pressure recordings were assessed at the level of the right carotid artery. All the analyses were performed in LabChartPro (v8.1.19) following the PV-loop workflow. Pressure calibration was performed before each recording and 10mmHg was then added to the channel to correct for a systematic error.

### 2.5 Histochemistry

Hearts were excised, rinsed in PBS, quickly blotted on gauze, and then fixed in 10% formalin for 24 hr. The biventricular apex of the heart was embedded in paraffin and cut in 4 μm sections. Sections were stained with Masson’s trichrome (Polysciences, Inc., Warrington, PA, USA) to assess collagen abundance. Stained sections were scanned (20x magnification) with AxioScan Z1 (Carl Zeiss, Jena, Germany), to obtain whole cross-sections for collagen quantification. Total fibrosis area (%) and perivascular fibrosis (ratio of the area of fibrosis surrounding the vessel wall to the lumen area) were quantified using ZEN2 blue edition (Carl Zeiss). All histological quantifications were independently performed by trained researchers blinded to groups.

### 2.6 Samples for spectroscopic analyses

Snap-frozen sections of murine cardiac ventricles were used. Ventricular sections of a total of six replicas of Ln and Ob tissues were analyzed by Raman and FTIR spectroscopies. In addition, a tissue-print procedure was applied to each sample to further extend the analysis of these non-fixed specimens. In particular, RV and LV sections were impressed on CaF_2_ windows to get the deposition of thin films of the extracellular matrix that were inspected by FTIR in transmission mode. Since the heart tissue is homogeneous on the spatial scale of the resolution achievable with vibrational spectroscopic techniques, mean spectra characteristic of different types of tissues and films (Ln and Ob) were compared.

### 2.7 Raman measurements

Raman scattering spectra were recorded with an Olympus IX73 inverted confocal microscope coupled to the S&I MonoVista CRS+ spectrometer, directly on snap-frozen tissue sections of cardiac ventricles. An excitation wavelength of 785 nm, with a power of 13.6 mW, was used in backscattering mode, along with a grating of 300 groves/mm and a 10x objective lens (N.A. = 0. 30). Each spectrum was the average of 60 accumulations at 10s integration time in the 400 cm^-1^ – 1942 cm^-1^ range, with a spectral resolution of ~2 cm^-1^. Spike removal was performed by Origin 2022 software from OriginLab.

### 2.8 FTIR-Attenuated Total Reflection (ATR) measurements of non-fixed cardiac tissue

FTIR-ATR spectra were collected directly on the LV and RV of heart tissues by using a Bruker Tensor 27 spectrometer and a Hyperion 3000 microscope equipped with a 20× ATR-objective, which is provided with a germanium crystal, and a 64 × 64 pixel nitrogen-cooled focal-plane-array detector. The 900 cm^-1^ – 3800 cm^-1^ spectral range was collected with a spectral resolution of 6 cm^-1^ and 512 scans. FTIR images (4096 spectra for each map) were collected in two different regions of each ventricle and corrected with the proper atmospheric compensation routine of OPUS 8.1 from Bruker Optics.

### 2.9 FTIR measurements of tissue print (ECM film)

ECM films were analyzed in transmission mode by FTIR spectroscopy. Spectra were collected using a Bruker Tensor 27 spectrometer and a Hyperion 3000 microscope equipped with a 15x Cassegrain objective and a nitrogen-cooled single-element mercury cadmium telluride detector. For every sample, 75 randomly selected punctual measurements were performed and the spectral range from 750 cm^-1^ to 7500 cm^-1^, with a spectral resolution of 6 cm^-1^ and 512 scans, was recorded. All the spectra were corrected with the atmospheric compensation routine of OPUS 8.1, and a Standard Normal Variate Normalization in the range 900-1800 cm^-1^ was applied. FTIR spectra were analyzed as second derivative profiles.

### 2.10 Statistical analysis

Raman spectra were obtained from 20 different points of each tissue sample. All the following spectral manipulations were conducted in R-studio. A quality test was carried out to identify and discard poor quality data: spectra selection was done by setting a minimum threshold equal to 1/10 of the maximum integral intensity of the amide I band. A baseline correction was carried out by means of a fifth-order polynomial method. A frequency calibration was performed, setting the intense phenylalanine peak at 1003 cm^-1^. Spectra were normalized to the intensity of the same 1003 cm^-1^ peak area. To test the reproducibility of data, a Principal Component Analysis (PCA) was applied. A single PC does not necessarily represent a single molecular species, since different types of spectra can be distinguished by the variation of several bands with respect to others. The molecular species constituting the tissue were assigned to these bands and the dominant PC1, PC2 and PC3 axes were considered to identify outliers. The two groups of cardiac tissues from Ln and Ob rats are well discriminated using this method. Once the outliers were discarded from the dataset, at least 60 Raman acquisitions for each type of tissue, RV and LV from Ln and Ob rats, were averaged to obtain the mean Raman spectrum with corresponding standard deviation.

FTIR images were collected in two different regions of each ventricle. These regions were randomly selected over the entire sample area in order to obtain a spectrum that is representative of the whole sample. A quality test was carried out for the FTIR spectra to remove spectra with a low signal-to-noise ratio. The absorbance value of the amide I band (1620-1690 cm^-1^) was considered as the reference signal, while the absorbance values in the signal-free region (1800-1900 cm^-1^) as the noise. Spectra were selected by setting a minimum threshold equal to 1/10 of the maximum integral intensity of the amide I band and a signal-to-noise ratio higher than 80. Data were analyzed as second-derivative profiles. A Standard Normal Variate Normalization in the range 900-1800 cm^-1^ was executed to normalize spectra. As well as with Raman spectra, a mean (second derivative) profile and corresponding standard deviation was obtained for each sample type (LV and RV from both Ln and Ob rats).

The difference profiles obtained from Raman spectra were obtained by subtracting the Ln from the Ob data: in this way, we obtained positive features for those signals with a higher intensity in Ob samples. For IR measurements (both ATR-FTIR of tissues and transmission spectra of ECM films), the difference was evaluated by subtracting the Ob from the Ln spectra. In the second derivative spectra, the negative sign of IR spectral components determined the different choice, i.e. opposite sign in the difference. In this way, if the Ob-Ln difference of Raman spectra is compared to the Ln-Ob difference of IR second derivative profiles, the same interpretation can be adopted: a positive feature is related to an increase of intensity in Ob with respect to the Ln counterpart.

For the interpretation of the spectra, we referred to ^7,14–16^.

## 3. Results

### 3.1 ZSF1 rats: functional and structural assessment of the heart

Obese (Ob) rats exhibited a remarkable increase in body weight over their lean (Ln) counterparts (Figure S1a). Obesity was accompanied by elevated levels of circulating glucose, triglycerides, cholesterol, lipids (Figures S1b-e), as well as elevated blood pressure (Figure S1f).

In Ob rats, cardiac hypertrophy was evidenced by a 77% increase in heart weight (Figure S2a). Interstitial heart fibrosis in both LV and right ventricle (RV) was slightly elevated in Ob compared to Ln rats (Figures S2b,c). Perivascular collagen area to lumen area (PVCA/LA) appeared to be similar between Ln and Ob rats (Figure S2d).

### 3.2. Spectroscopy

#### 3.2.1. Principal Component Analysis of Raman data

RV and LV tissues taken from Ln and Ob animals were studied. A PCA was executed to better identify which spectral features were involved in the differentiation between tissues from Ln and Ob rats (a detailed description of these principal components can be found in the Supplementary Materials, SM, file). The score plots (Figures 1a,b) showed that, for both RV and LV, the two populations are clearly separated by the three main principal components, which describe more than 73% of the variance (Figure 1c) in the dataset.

**Figure 1.**
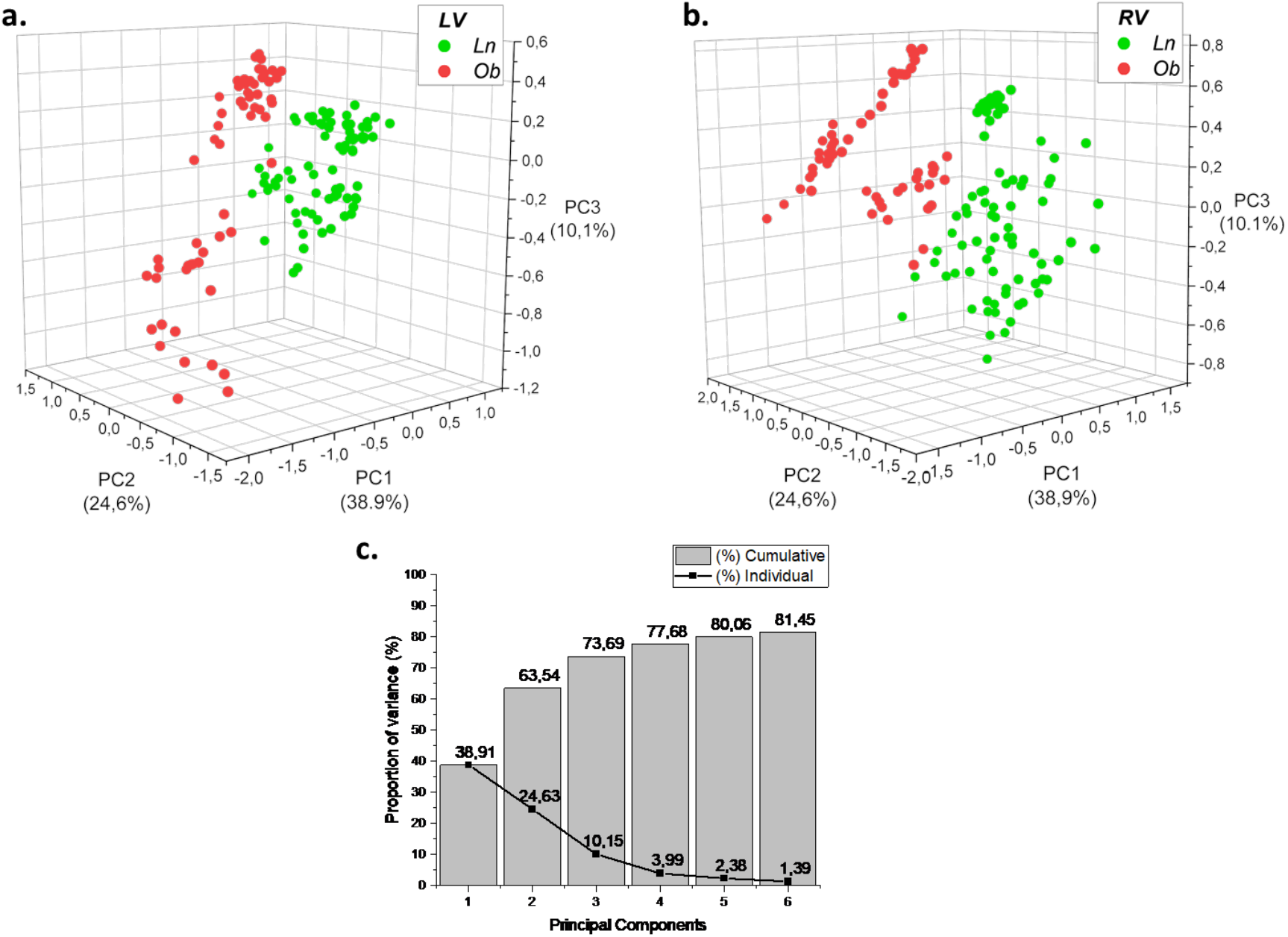
Principal component analysis of Raman data. Scores of PC1, PC2 and PC3 for the left (a) and right (b) ventricles. Cumulative and individual variances of the first six components of PCA (c).

In the case of the LV, the separation is entirely dependent on PC1. Specifically, PC1 loading (Figure S3a) shows positive contributions at 1238 and 1577 cm^-1^, and negative ones at 850, 932, 1046, 1300, 1451 and cm^-1^. Ln and Ob samples are grouped at positive and negative PC1 scores, respectively. PC2 represents more than 22% of the variance of the spectral dataset and it is characterized by predominantly negative contributions at 1259, 1340, 1441, 1605, and 1655 cm^-1^ (Figure S3b). Differently from the LV, the PCA analysis on spectra of the RV showed that Ln and Ob keep clustering up to PC3. Specifically, Ob samples have positive scores, whereas Ln scores are almost exclusively negative. PC3 loading shows negative contributions mainly at 1178 cm^-1^, while positive ones are at 841, 915, 971, 1049, 1229 and 1458 cm^-1^ (Figure S3c). The combination of the spectral contributions of PC1, PC2, and PC3 to the clear separation of Ln from Ob suggests the development of different modifications of the biochemical composition of RV compared to LV.

#### 3.2.2. Mean Raman spectra and different profiles of heart tissue

Data selected by PCA (a few outliers were eliminated) were averaged to obtain mean spectral profiles representative of LV (Figure 2a) and RV (Figure 2b) of Ln and Ob samples. To better highlight variations of spectral profiles, we also evaluated the difference spectrum, obtained by subtracting the mean spectrum of the Ln from the one of the Ob samples. The results are shown in Figures 2c and 2d. The positive contributions to the difference spectrum identify the wave numbers at which the signal intensity is higher in the Ob samples than in the Ln ones. Negative contributions, on the other hand, are found in bands where the intensity is higher in Ln samples than in Ob ones (see MM 4.10). Difference profiles show that the most significant variations in signal intensity from spectra of Ln to Ob samples are positive. In particular, we can identify three spectral regions in which positive contributions appear for both LV and RV tissue spectra: 830-950 cm^-1^; 1030-1100 cm^-1^; and 1420-1480 cm^-1^. A positive narrow contribution centered at 1660 cm^-1^ is particularly intense for the different profile of the LV (Figure 2c). Negative contributions are observed in the 1150-1260 cm^-1^ and 1500-1625 cm^-1^ spectral ranges (Figure 2c,d).

**Figure 2.**
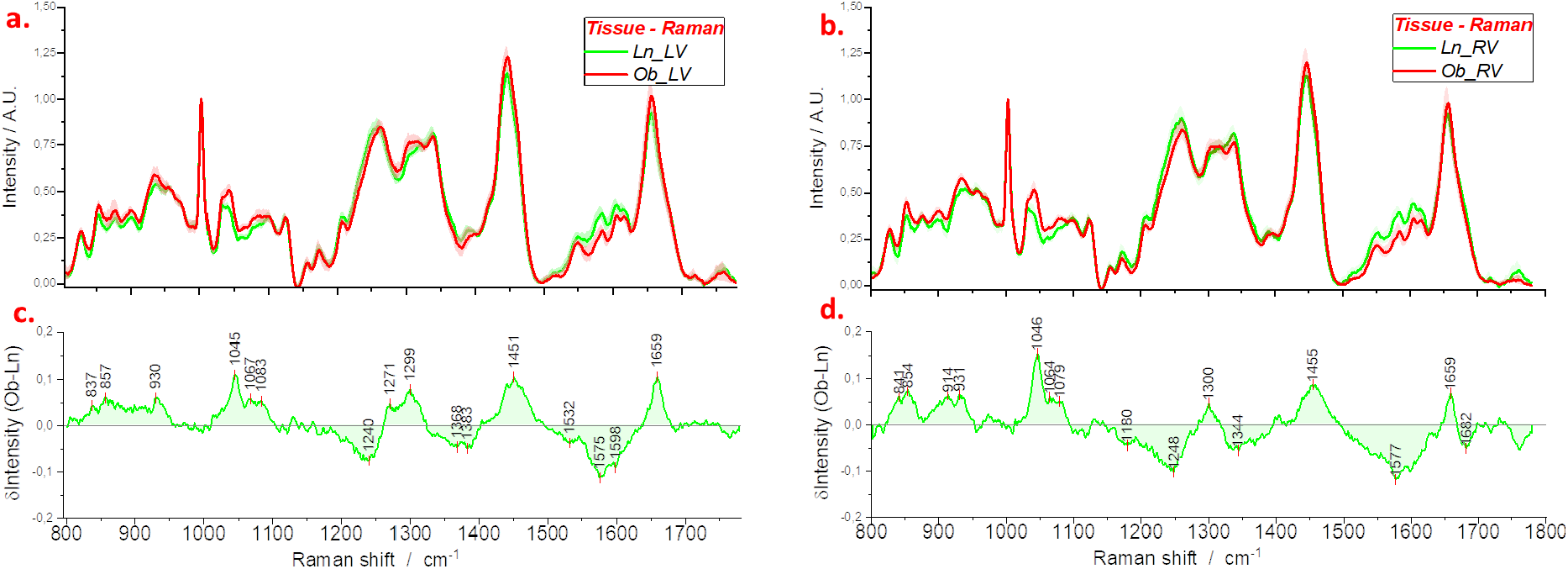
Mean Raman spectra obtained from tissues of the left (a) and right (b) ventricles of Ln and Ob rats; standard deviations are reproduced by shadowed areas. Difference profiles in panels (c) and (d) are obtained by subtraction of the green from the red curve of panels (a) and (b), respectively.

#### 3.2.3. Mean Attenuated Total Reflection (ATR)-FTIR spectra and different profiles of heart tissues

To further inspect the average composition of Ln and Ob tissues, the same samples investigated by Raman spectroscopy were analyzed by ATR-FTIR. The average ATR spectra of LV and RV samples obtained from Ln and Ob groups are shown in Figure 3 as second derivative spectra. In this representation, the intensity of a spectral component is more and more negative when increasing the concentration of the molecular species assigned to the band. Figures 3a and 3b show both data groups with their standard deviations. Intensity variations from Ln to Ob are hardly detected; the few significant changes are larger for the RV and particularly in the 1480-1600 cm^-1^ spectral range. This is better evidenced in Figures 3c and 3d: in these panels, the difference between the second derivative profiles of Ln and Ob samples is shown. An increase in the intensity of specific components of the Ob spectra, compared to Ln spectra, is related to positive peaks of the difference (see Figure S3). Larger increases in the intensity for Ob samples are evidenced at 1560, 1740 and 2930 cm^-1^ for RV, and at 1645, 1740 and 2930 cm^-1^ for LV. The negative peak at 1506 cm^-1^ in the difference profile of RV (Figure 3d) indicates a decrease of intensity for this spectral component passing from Ln to Ob spectra.

**Figure 3.**
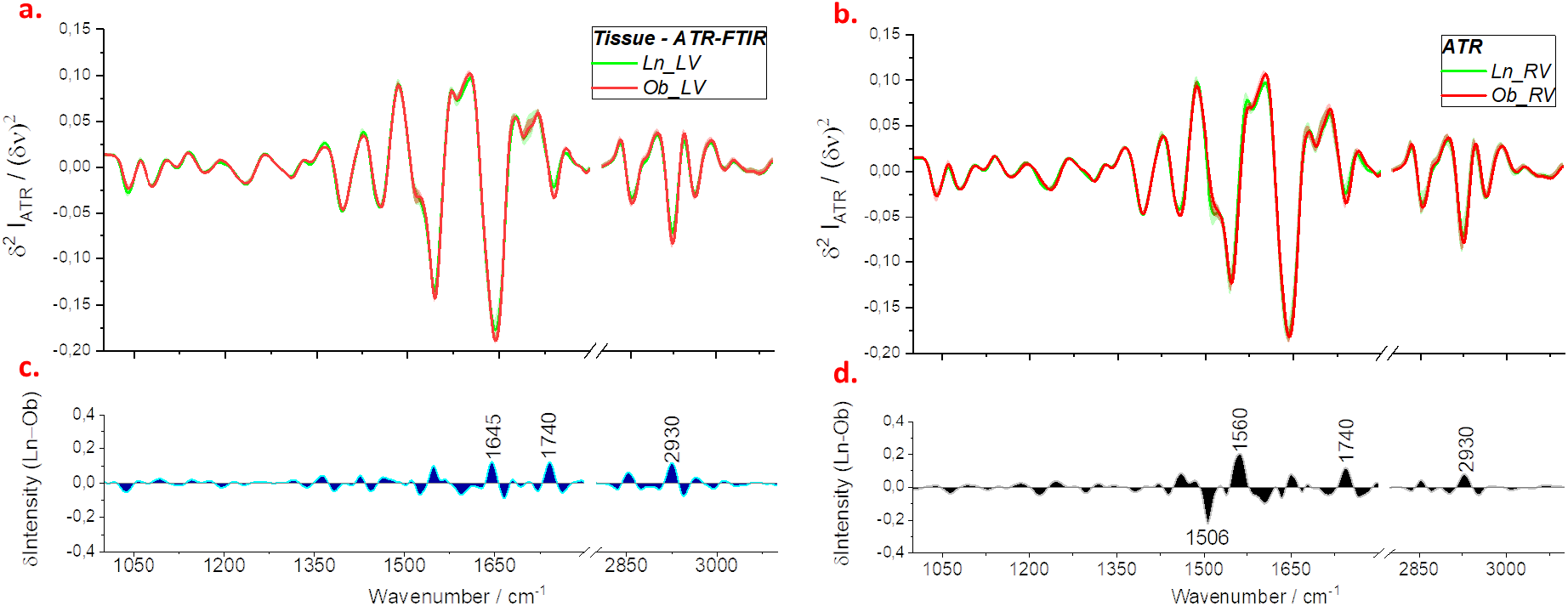
Mean ATR-FTIR spectra obtained from tissues of the left (a) and right (b) ventricles of Ln and Ob rats. Difference profiles in panels (c) and (d) are obtained by subtraction of the green from the red curve of panels (a) and (b), respectively.

#### 3.2.4. Mean FTIR spectra and different profiles of ECM films

The ECM was further investigated by imprinting the surface of the fresh section on a CaF2 window to achieve the IR spectrum in transmission mode. In Figure 4, IR measurements of tissue prints are presented. The different spectral components are recognized on the negative intensity of the second derivative spectra shown in Figures 4a and 4b; the intensity variation is shown by the difference profiles presented in Figures 4c and 4d. The increase or decrease in concentration for a molecular species of Ob compared to Ln spectra is related to the positive or negative peak of difference profiles, respectively (see SM). When the whole tissue and ECM assessments are compared, it is remarkable to see that differences are larger for spectra from ECM, and this is predominant for the RV in the regions 2800-2950 cm^-1^ and 1600-1800 cm^-1^. The only exceptions are the peaks at 1506 and 1560 cm^-1^: the intensity variation at these frequencies is larger for the different profiles of the whole tissue rather than ECM film.

**Figure 4.**
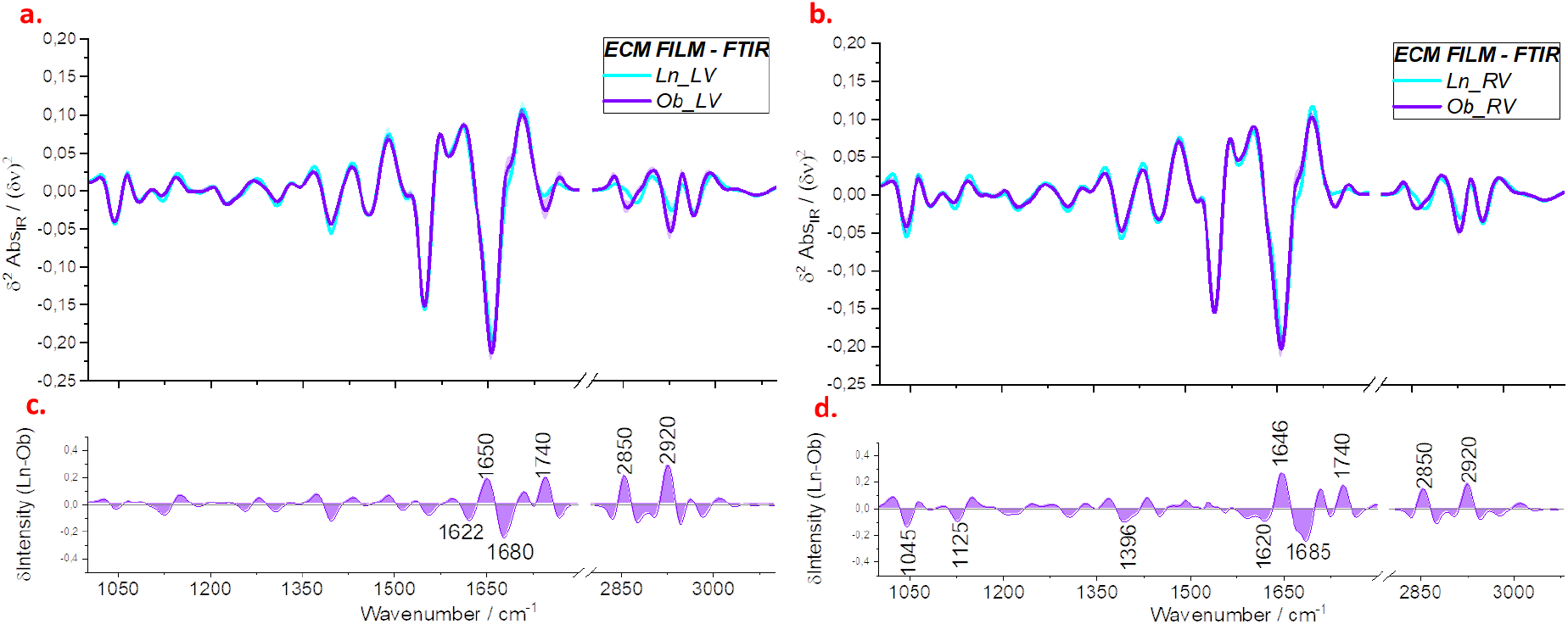
Mean FTIR spectra obtained from tissue prints of the left (a) and right (b) ventricles of Ln and Ob rats. Difference profiles in panels (c) and (d) are obtained by subtraction of the violet from the blue curve of panels (a) and (b), respectively.

## 4. Discussion

ZSF1-Ob rats presented characteristic signs of adverse cardiac remodeling and function. Ob rats exhibited concomitant elevated levels of lipids, triglycerides, glucose, and high blood pressure. Also consistent with the HFpEF phenotype, the Ob rats showed no evidence of compromised systolic function or ventricular dilation, elevated cardiac stiffness and increased interstitial ventricular fibrosis.

In the analysis of cardiac sections, Raman and ATR data showed a different sensitivity toward molecular components of the tissue, though the two techniques both monitor the average chemical composition of the heart. Spectroscopic data suggested that HFpEF is associated with a sensitive change in tissue biochemistry since peaks assigned to cellular and extracellular matrix demonstrated a significant variation of intensity in Ob spectra with respect to samples from Ln rats. Both FTIR and Raman spectra indicated an increase of lipid concentration in the tissues of Ob rats: this was monitored by the Raman bands at 1300 and 1450 cm^-1^, and by the IR signals at 1740, 2850 and 2930 cm^-1^. The enhancement of these bands was almost the same in the spectra from the two ventricles suggesting that this effect is similar for the two chambers. In addition, the increase in lipid concentration was particularly evident in the ECM of both ventricles.

In HFpEF, diastolic dysfunction is often observed due to stiffness of the myocardium and changes in the ECM. The variation of Raman profiles in the regions of amide I (1625-1700 cm^-1^) and amide III (1200-1300 cm^-1^) bands of Ob samples are compatible with a rearrangement of collagen fibers and particularly with an increase of order in the alignment of bundles. This was indicated by the decrease of 1245 cm^-1^ intensity and the corresponding enhancement of 1660 cm^-1^ spectral component^17^. A different result was obtained for the heart tissue of a different animal model of HFpEF, induced by a high salt diet^4^. On tissues from Dsalt-sensitive (DSS) rats, an increase of intensity for the amide III band, together with a blue shift of the amide I band was assigned to the increase of collagen concentration in diseased compared to normal animals. In this experiment, rather than an increase in collagen concentration, we observed a structural change of collagen fibers.

Experimental evidence reported in previous studies has shown that the stiffening of cardiac tissue can be related to the altered organization of the collagen network^18^. It was observed that augmented myocardial stiffness is accompanied by structural rather than concentration variations in total myocardial collagen or collagen phenotypes^18^. Abnormal cross-linking between collagen fibers is involved in these structural modifications of the network, which has been referred to as a major determinant of LV diastolic dysfunction in patients with HF. From a pathophysiological point of view, high myocardial collagen cross-linking (CCL) may reduce diastolic reserve and thus, in conditions of fluid overload or exercise, the subsequent acute elevation of filling pressures may lead to reduced diastolic filling and produce HF symptoms^19^.

In the Raman spectrum, a possible marker for the presence of CCL is the frequency shift of the amide III band. The increase in cross-linking mechanisms causes collagen fibers to be more tightly connected to each other. This collagen network tightening may limit the rotational freedom of individual fibers, leading to an increase in ordered structures. This effect could explain the narrowing of amide I peak in the IR profile of ECM film, that is evidenced in the difference profiles shown in Figures 4c and 4d by the two negative contributions on both high (ca. 1680 cm^-1^) and low (ca. 1620 cm^-1^) frequency sides of amide I maximum at about 1650 cm^-1^.

According to histochemical analysis (Figure S2), interstitial heart fibrosis of ZSF1 rats is increased in LV and RV of Ob in comparison to Ln (Figures S2c,d). Perivascular fibrosis appears to be low and similar between Ln and Ob rats (Figure S2e). Differently, in DSS rats^4^ abundant collagen deposition was observed in both perivascular and interstitial areas. Thus, the differences between spectroscopic data obtained for the two models, are coherent with histochemical results.

A pronounced intensity variation between spectra of healthy and diseased samples is found in the 1550-1620 cm^-1^ region of the Raman spectrum. This region is characteristic of vibrations of the aromatic amino acids such as phenylalanine (Phe), tyrosine (Tyr), and tryptophan (Trp), but only Trp shows an intense Raman band at about 1580 cm^-1 20^. The significant intensity change at this frequency may therefore be associated with the specific decrease in Trp content in the Ob tissue. This finding provides important information about the inflammatory state of the diseased tissue, which is a favorable condition for the development of HF. In fact, it was observed that the increase in cytokine production induced by the inflammation state is accompanied by a decrease in Trp^21^.

The reduction in Trp content may also be related to a different mechanism affecting the stiffness of the tissue cellular component. Trp is an essential component of Titin protein which plays a key role in the contraction and relaxation of cardiomyocytes^22^. A reduction of Trp may result in altered structure and function of Titin leading to increased stiffness^23–25^. We hypothesized that a sign of this effect can be recognized in the band intensity decrease in the 1550-1620 cm^-1^ region of Raman spectra and at 1506 cm^-1^ of the ATR-FTIR spectra, associated with Trp, in the tissue of diseased rats.

In ATR, an increase of intensity is observed in the spectral profile of Ob compared to Ln at ~1560 cm^-1^, the frequency at which a vibration from glutamate can be observed. The increase in glutamate concentration can be regarded as a marker of HF onset, according to the study by Ariyoshi M. et al. ^26^. The mitochondria of cardiac cells contain the protein 9030617O03Rik, whose expression drops significantly in the presence of HF. The reduction of 9030617O03Rik was noted to induce a significant increase in the concentration of glutamate in the heart^26^. The fact that both the augmentation of glutamate and the decrease of Trp concentration relate to the different activity of the cellular constituents of the tissue is confirmed by the absence of relevant intensity variations in the spectra from ECM films.

A difference in the metabolism of Ob vs Ln cells is also indicated by the Raman bands in the region 1000-1100 cm^-1^, which are characteristic of carbohydrates or glycosidic units. Cellular dysfunction underlies the pathogenesis of the obesity-associated metabolic disease, including type 2 diabetes. Thus, the presence of high carbohydrate concentration in the Ob cardiac tissue is not surprising. Interestingly, this abnormal carbohydrate concentration is also found in the ECM and this is evidenced by the IR data shown in Figure 4d.

## 5. Conclusions

This work demonstrated the potential of vibrational spectroscopic techniques to monitor the biochemical changes present in HFpEF. These spectroscopic tools proved to be very sensitive in detecting changes in the cardiac chemical composition, which may contribute to the alteration in the mechanical properties of this cardiac disease.

Comparison of spectral profiles, particularly Raman spectra, between normal and diseased samples allowed reconstruction of the possible structural and biochemical changes that can induce the tissue stiffening. It is reported in the literature that cardiac tissue stiffening is due to variations in both the extracellular and cellular components; in ECM, in particular, a fundamental role is played by collagen. For the specific analysis of ECM, we used tissue printing as a novel analytical tool. The tissue printing method enabled the analysis of ECM as well as the production of replicas of the tissue to perform different analyses. Since these prints can be stored at low temperatures and analyzed later, this approach is especially useful with non-fixed tissue samples.

According to our data, the cardiac tissue of Ob rats does not exhibit an important collagen deposition or type III to type I phenotypic shifts but rather abnormal cross-linking causing the stiffening of ventricles. The stiffening is also evidenced by the decrease of Trp related to the altered Titin structure, and to the inflammatory state which is a favorable condition for HFpEF.

The altered biochemistry of cardiac cells was also recognized in the augmentation of glutamate and glycoproteins monitored in both ventricles but particularly in the right chamber. This asymmetry of altered biochemistry in the two ventricles was observed by IR and Raman analyses of tissues also in a different model of HFpEF ^4^. However, in the case of hypertension induced by a high-salt diet, the late stage of HFpEF correlated to the formation of micro-calcifications, which were not observed in the present work. Thus, our study suggests that comorbidities play a fundamental role in organ damage, and a sensitive and selective probe of the tissue biochemistry is fundamental to provide for a specific therapeutic treatment. Overall, these results suggest the presence of specific chemical fingerprints in this model of HFpEF, with increased glutamate and glycoproteins, increased lipids, formation of cross-linked collagen, and decreased Trp content. This differs from the structural change of tyrosine units, the free amino-acid and collagen accumulation, and the formation of cardiac micro-calcifications that have been observed in DSS rats and are perhaps specific to hypertension-induced HF. These findings could aid in better defining the pathological mechanisms underlying different forms of HF. The clinical applicability of these observations will be further increased when the corresponding circulatory fingerprints of these cardiac chemical modifications are determined. Future studies are necessary to identify such early, novel circulatory markers of these spectroscopically detected chemical modifications in cardiac tissue.

## Supplementary Materials

Figure S1. Cardiac function and biochemistry: Figure S2. Cardiac structure and histology; Figure S3. Loadings from PCA analysis of Raman data.

## Author Contributions

Conceptualization, A.C., P.S.; methodology, A.C., P.S. and L.P.; formal analysis, L.P., data curation, L.P.; writing—original draft preparation, M.P., L.P. and P.S.; writing—review and editing, P.S. and A.C., R.P., A.S., GJJ.S.; supervision, P.S. and A.C.; funding acquisition, P.S. and A.C. All authors have read and agreed to the published version of the manuscript.

## Acknowledgments

The authors thank Niki Tombolesi for his support in defining the computational protocol for the manipulation of spectroscopic data, and Raffaele Altara for the careful handling of the animals.

## Supplementary Materials

**Figure S1.**
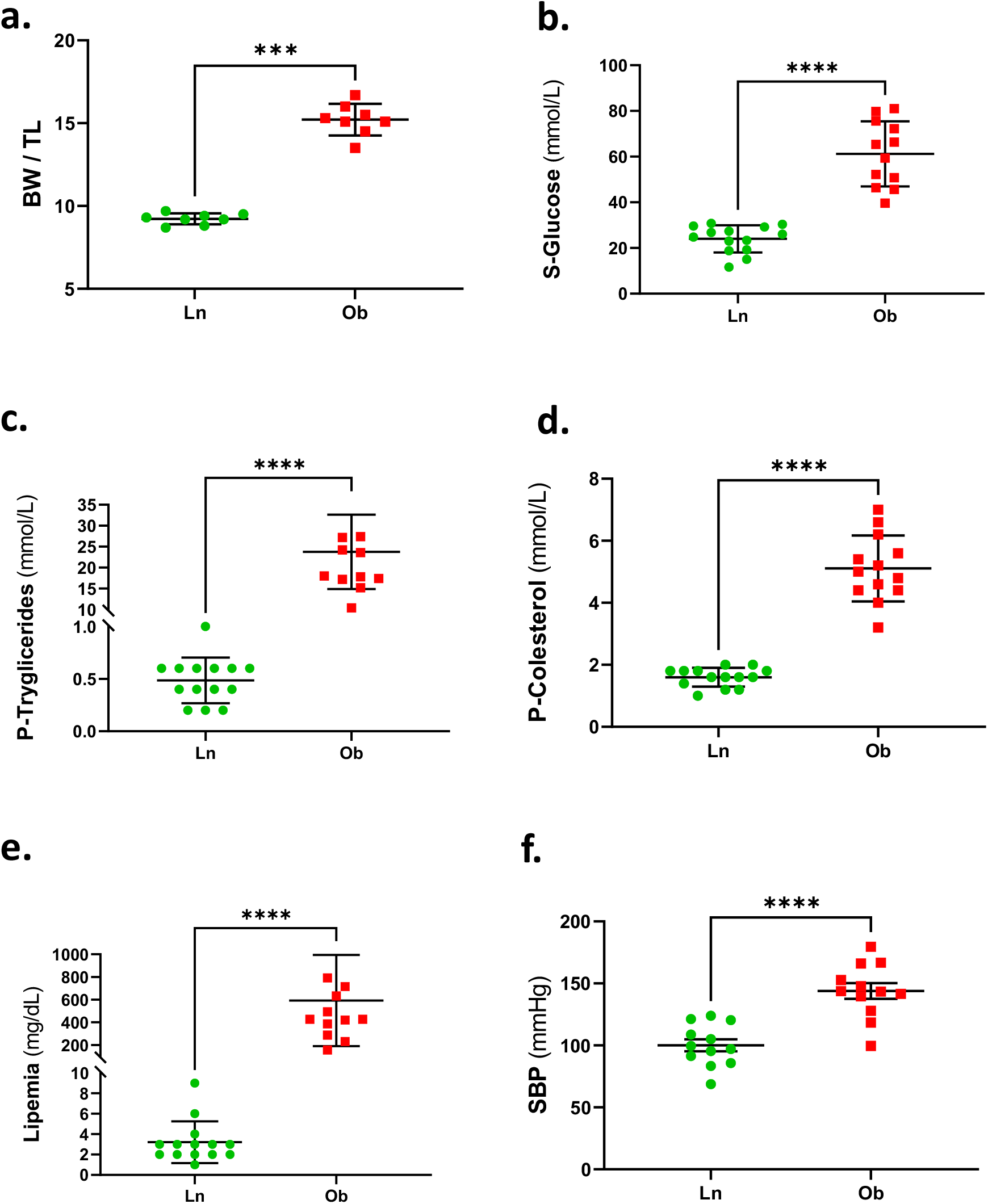
Cardiac function and biochemistry: body weight (BW); levels of circulating glucose (S-Glucose), triglycerides (P-Tryglicerides), cholesterol (P-Cholesterol), and lipids (lipidemia) in blood serum; systolic blood pressure (SBP).

**Figure S2.**
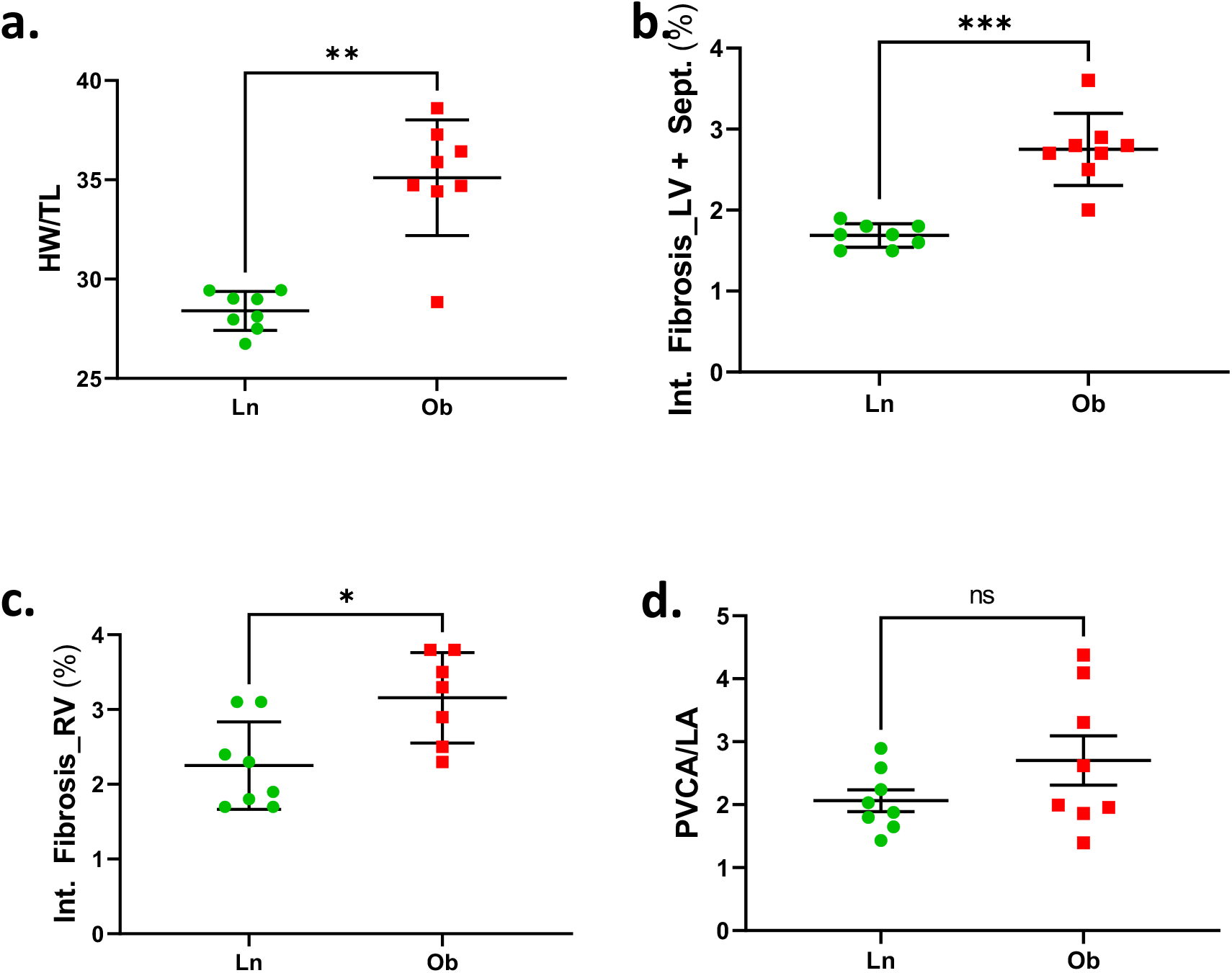
Cardiac structure and histology: heart weight (HW); interstitial fibrosis (Int. Fibrosis) of left (LV) and right (RV) ventricles; Perivascular collagen area to lumen area (PVCA/LA).

**Figure S3.**
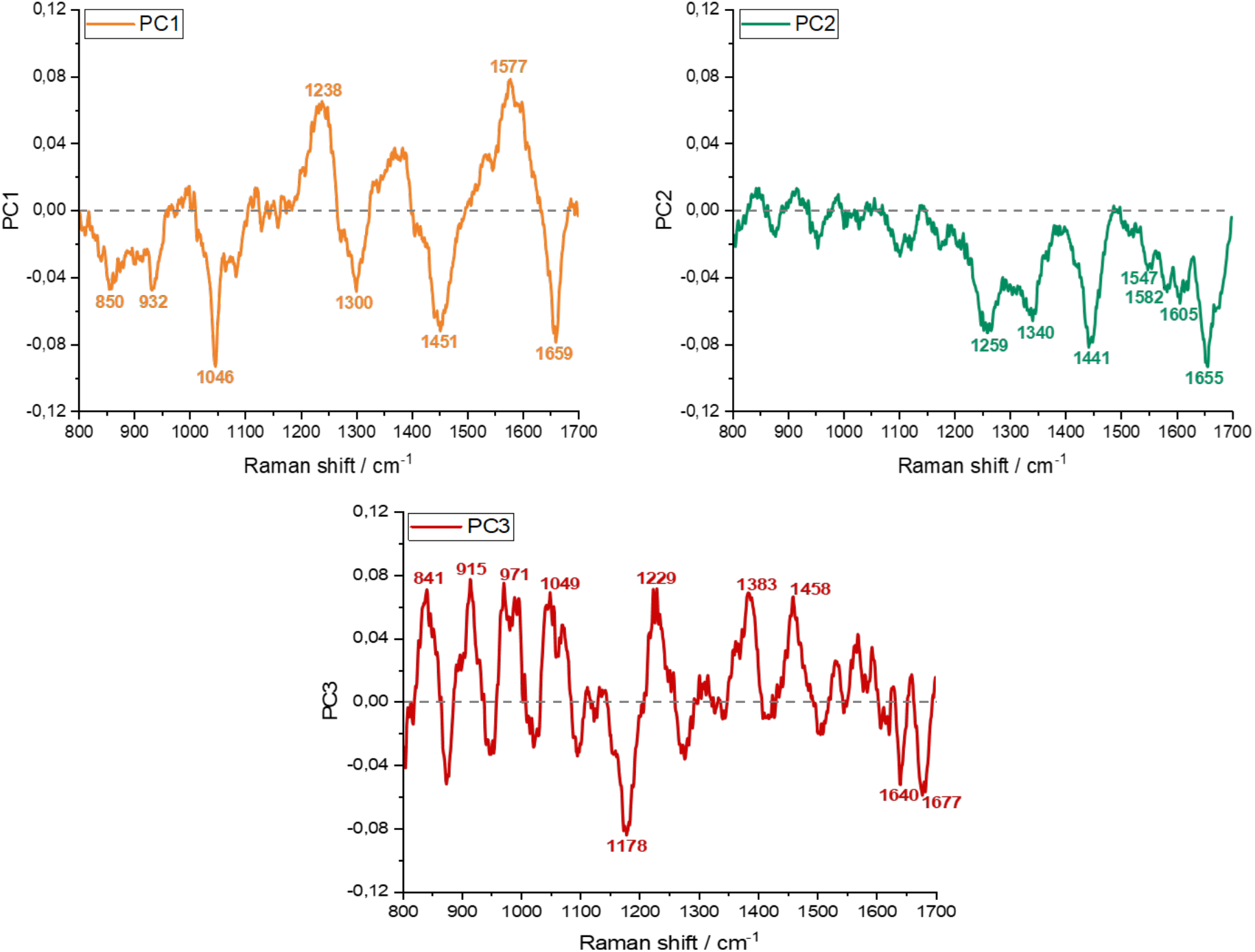
Loadings from PCA analysis of Raman data. Principal Component Analysis is a commonly used multivariate approach able to reduce the dimensionality of large datasets. It can be applied to spectroscopic data to identify the most variable spectral features of a sample (or sample’s area) based on differences in biochemical content. In this approach, loadings represent spectral variations which differentiate the groupings according to the intensity measured at each wavenumber. The two groups of cardiac tissues from Ln and Ob rats are well discriminated using this method, and the basis for discrimination is derived from analysis of the loadings corresponding to each PC. The minima and maxima of each PC are analyzed in order to recognize the spectral components (bands) of the pristine spectrum; differently from spectra, the bands of loadings can be either positive or negative. The molecular species constituting the tissue are assigned to these components. A single PC does not necessarily represent a single molecular species, since different types of spectra can be distinguished by the variation of several bands with respect to others. These means that different types of tissue can be distinguished because more than one molecular species vary with respect to others. The PC1 loading reported above shows the signals characteristic of tryptophane (1577 cm^-1^), proteins (930, 1238, 1300 and 1659 cm^-1^), lipids (1451 cm^-1^) and glycoproteins (1046 cm^-1^). The opposite sign of 1577 and 1238 cm^-1^ features with respect to others suggests that Trp (1577 cm^-1^) and less ordered structures of collagen (1238 cm^-1^) show an opposite variation with respect to the other molecular species (highly ordered collagen fibers, lipids and glycoproteins). In Ob-LV samples, the two features decrease, and the others increase in comparison to Ln-LV samples. The clustering of RV tissues is also depending on the intensity change at 1178 cm^-1^(intense negative feature of PC3) which is assigned to proteins.

